# Nanocluster-Mediated Signaling Crosstalk between FcγR and TLR4 in Macrophage Inflammatory Responses

**DOI:** 10.1101/2024.09.27.615544

**Authors:** Seonik Lee, Yan Yu

## Abstract

Receptor crosstalk, the interaction between different receptors to modulate signaling, is crucial for fine-tuning the inflammatory responses of innate immune cells. Although the synergistic crosstalk between Toll-like receptor (TLR)4 and Fc gamma receptor (FcγR) is well documented, the detailed mechanism underlying this synergy remains unclear. In this study, we addressed this knowledge gap by imaging the molecular organization of TLR4 and FcγR on the macrophage cell surface and correlating it with their synergistic co-activation using ligands functionalized on lipid bilayers. We confirmed that co-activation of TLR4 and FcγR enhances whole-cell pro-inflammatory responses and tyrosine phosphorylation at the receptor level. Super-resolution microscopy revealed that TLR4 and FcγR each form discrete nanoclusters after ligand stimulation, and their synergistic co-activation significantly increases both the size and spatial overlap of these nanoclusters, leading to amplified immune signaling. Contrary to previous assumptions that TLR4 and FcγR form heterodimers during their crosstalk, our results emphasize the critical role of receptor nanocluster interactions in modulating innate immune responses. Additionally, these findings align with similar receptor interaction mechanisms that we previously reported in other receptor pairs, such as Dectin-1/TLR2 and FcγR/TLR2, suggesting that nanocluster interactions may represent a predominant mechanism governing crosstalk between TLRs and ITAM-containing receptors.

## Introduction

Innate immune cells, including macrophages and neutrophils, are responsible for eliminating harmful microorganisms^1-3^. They utilize pattern recognition receptors (PRRs) to detect and respond to the so-called pathogen-associated molecular patterns (PAMPs). For example, the family of Toll-like receptors (TLRs) detect a diverse range of bacterial cell wall components such as lipopolysaccharides, while Fc gamma receptors (FcγRs) bind antibodies^2,4^. Although the functions of individual receptors have been extensively studied, much less is known about how different types of receptors coordinate immune responses. For instance, simultaneous stimulation of FcγRs with TLR2 or TLR4 has been reported to enhance the pro-inflammatory response of M2 macrophages in the context of rheumatoid arthritis^5^. Additionally, FcγR-TLR2 signaling crosstalk was found to synergistically regulate cytokine IL-6 production in murine macrophages^6^. In contrast, stimulation of the inhibitory FcγRIIb suppresses TLR4 responses in dendritic cells^7^. Similarly, activation of cytosolic RIG-I-like receptors downregulates gene transcription induced by TLR signaling^8^. While these observations highlight the prevalence of receptor signaling crosstalk, the mechanisms governing the interactions between different receptors during this process remain largely unknown.

The prevailing presumption has been that different types of receptors form heterodimers to enable their signaling crosstalk^9,10^. However, our recent research has uncovered a different mechanism involving interactions between nanoscale receptor clusters. With Dectin-1 and TLR2^11^, as well as FcγR and TLR2/1^12^, we have shown that each receptor forms its own discrete nanoclusters. It is the interaction between these individual receptor nanoclusters that enables the pairs of receptors to synergize their signaling. The crucial question that remains is: How broadly applicable is this mechanism to other pairs of immune receptors?

In this study, we investigate the spatial organization of FcγRs and TLR4 and how their interaction synergistically modulates proinflammatory responses in macrophages. Previous studies have suggested signaling crosstalk between FcγRs and TLR4 in innate immune cells. Specifically, the activation of TLR4, in conjunction with FcγR, increases cytokine production, thereby regulating inflammatory responses of THP-1 cells^13^. Observations from western blots and immunoprecipitation analyses have suggested an association between FcγRIII and TLR4, leading to speculation that these receptors form heterodimers to facilitate signaling crosstalk^9,10^ . However, no direct evidence has been provided to support the assumption of receptor heterodimerization.

Building on our previous findings that TLR2/Dectin-1 and TLR2/FcγR interact as nanoclusters to enable signaling crosstalk, we hypothesize that the nanoscale clustering of FcγRs and TLR4 similarly governs their signaling crosstalk. To test this hypothesis, we combined super-resolution fluorescence microscopy to quantify receptor clustering with biochemical assays to measure macrophage inflammatory responses. By using ligands conjugated on reconstituted lipid membranes with precisely controlled ligand densities, we confirmed that co-activation of FcγR and TLR4 enhances inflammatory responses in RAW264.7 macrophage cells, including receptor tyrosine phosphorylation, activation of transcriptional nuclear factor κ-light-chain-enhancer of activated B cells (NF-кB), and cytokine tumor necrosis factor (TNF)-α secretions. Super-resolution microscopy revealed that FcγR and TLR4 form discrete receptor nanoclusters of 50-100 nm, which increase in size and partially overlap during synergistic signaling crosstalk. Our findings provide direct evidence that the nanoscale clustering of FcγRs and TLR4 drives their synergistic regulation of macrophage inflammatory responses, challenging previous assumptions that these receptors form heterodimers during crosstalk.

## Results

### Functionalization and characterization of ligand functionalized lipid bilayers

In this study, we activated macrophage cells using ligand-functionalized planar lipid bilayers, which allowed feasible control over the conjugation density of the ligands – lipopolysaccharide (LPS) for TLR4 and immunoglobulin G (IgG) for Fc gamma receptor (FcγRs)^14,15^. RAW264.7 macrophage cells predominantly express FcγRI, FcγRIIB, FcγRIII, and FcγRIV, which bind to IgG^16^ . To co-activate both receptors, the lipid bilayers (referred to as LPS+IgG bilayers) were composed of 1,2-dioleoyl-sn-glycero-3-phosphocholine (DOPC), 2 mol% 1,2-dioleoyl-sn-glycero-3-phosphoethanolamine-N-(cap biotinyl) (biotin-DOPE) for tethering biotinylated IgG via streptavidin linkers, and 1 wt% of LPS (**Fig. 1a**). For selective activation of either TLR4 or FcγR, only LPS or IgG was included in the bilayers. By mixing fluorescent and non-fluorescent ligands and imaging them on lipid bilayers with total internal reflection fluorescence (TIRF) microscopy (**Fig.1b**), we estimated the ligand densities to be (4.6 ± 0.2) ×10^3^ IgG/µm^2^ and (1.7 ± 0.3) ×10^3^ LPS/µm^2^. To characterize the diffusion coefficients of the lipids and LPS, we incorporated 0.05 mol% Rhodamine-DOPE and 1 wt% Alexa488 dye-labeled LPS in the bilayers and used fluorescence recovery after photobleaching (FRAP) microscopy. The diffusion coefficients were 1.78 ± 0.05 µm^2^/s for lipids and 1.53 ± 0.10 µm^2^/s for LPS (**Fig. 1c, d**), indicating that LPS was successfully incorporated without impacting bilayer fluidity.

**Fig. 1.**
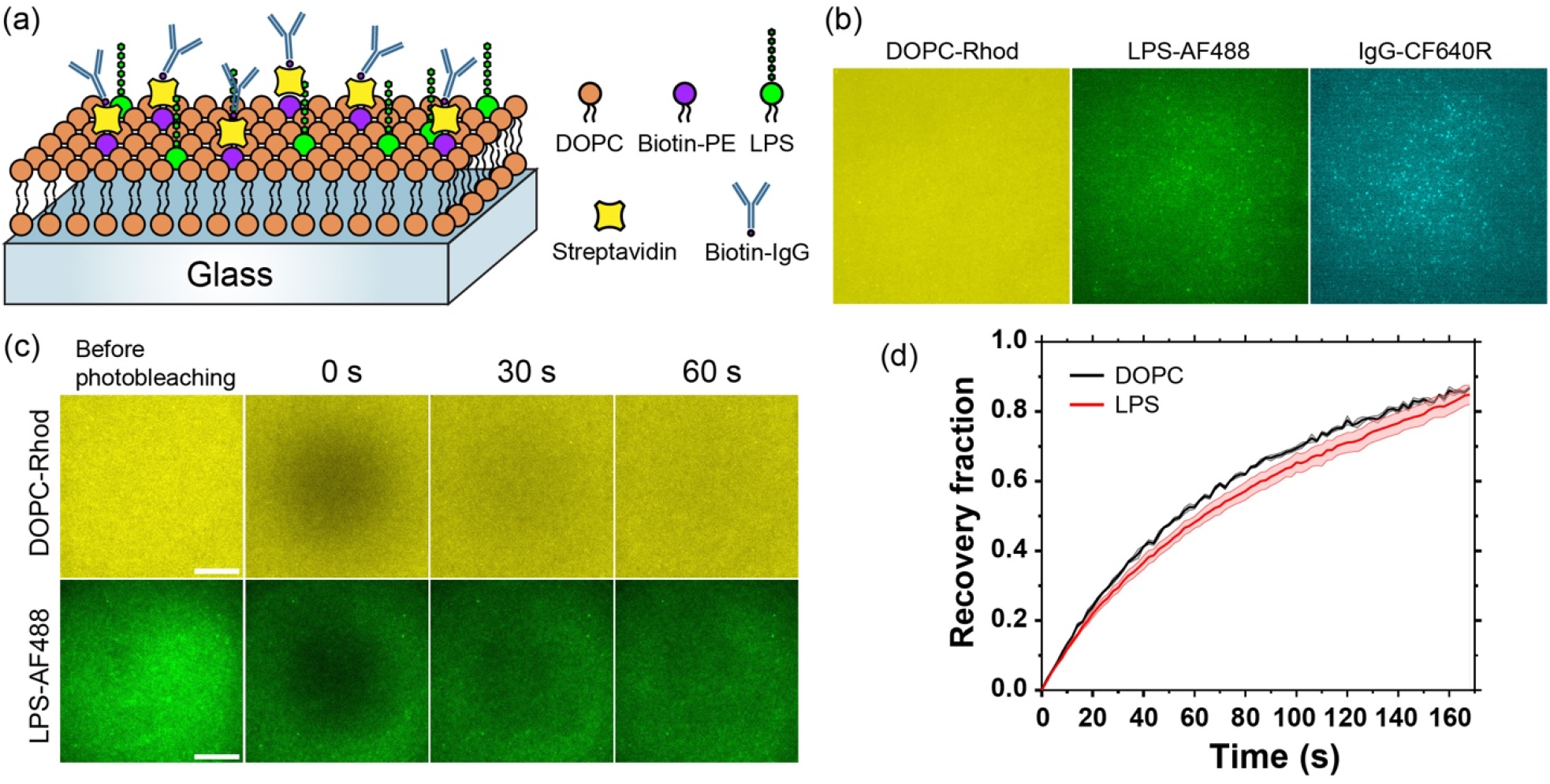
Ligand-functionalized lipid bilayers for activation of macrophage cells. (**a**) schematic illustration of the LPS+IgG bilayer. (**b**) Representative fluorescence images showing a lipid bilayer functionalized with Alexa Fluor 488-labeled LPS and CF640R-labeled IgG. Scale bar: 10 μm. (**c**) Fluorescence images showing the fluorescence intensity recovery after photobleaching (FRAP) of a lipid bilayer containing Rhodamine-DOPE (0.05 mol%) and Alexa Fluor 488-labeled LPS (1 wt%) at 37°C. Scale bars: 10 μm. (**d**) FRAP data of the lipid bilayer shown in (**c**). Each set of data is presented as mean ± SE (standard error) from three independent experiments.

### Co-activation of TLR4 and FcγR enhances cell pro-inflammatory responses

We next examined how FcγR and TLR4 crosstalk influences the pro-inflammatory response in macrophage cells, focusing on the translocation of nuclear factor κ-light-chain-enhancer of activated B cells (NF-κB) and the generation of inflammatory cytokine tumor necrosis factor (TNF)-α. NF-κB is a cytoplasmic transcription factor that translocates to the nucleus in response to external stimuli, such as pathogen recognition, where it regulates gene transcription and inflammatory cytokine production ^17,18^. We used RAW264.7 cells expressing RelA-EGFP, a subunit of the NF-κB family also known as transcription factor p65^19^, to directly image and quantify its cytoplasm-to-nucleus translocation (**Fig. 2**). The translocation of RelA-EGFP serves as a marker of canonical NF-κB signaling ^20,21^.

**Fig. 2.**
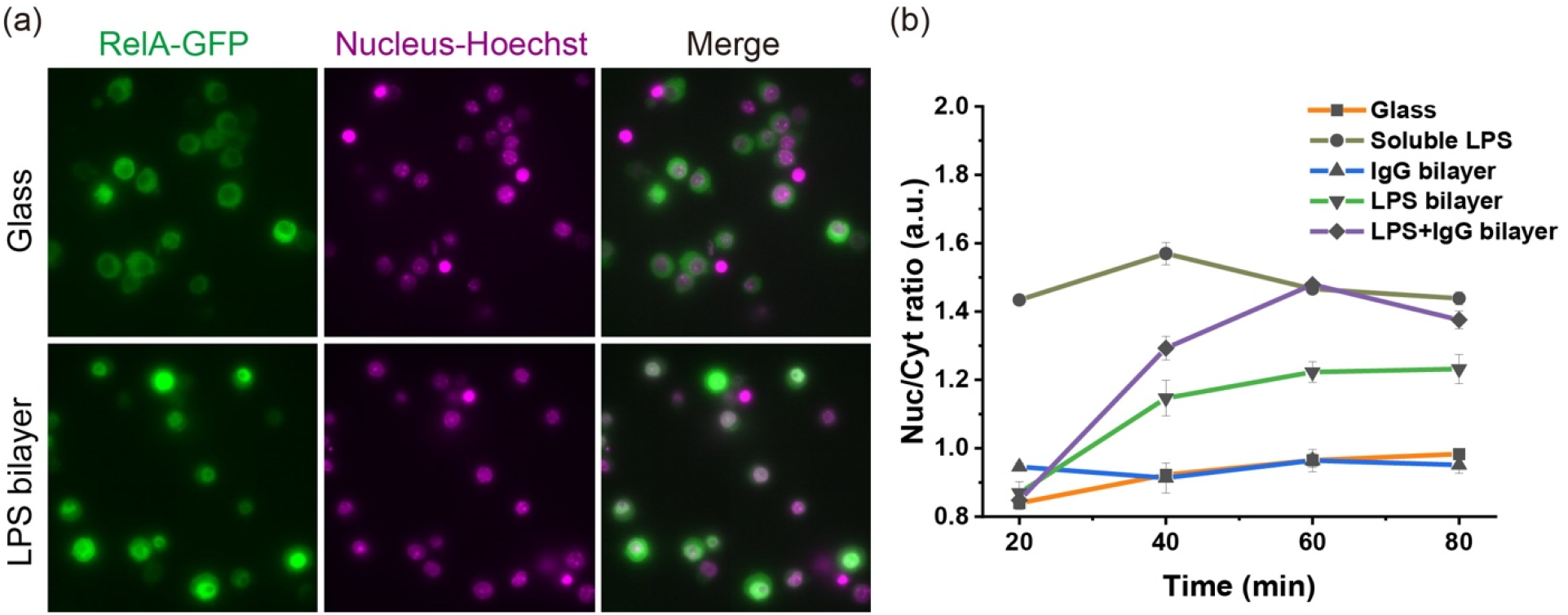
Quantification of NF-κB activation in RAW264.7 macrophage cells. (**a**) Epi-fluorescence images of cells stably expressing EGFP-RelA (green) and labeled with nuclear counterstain Hoechst (magenta) after 60 min incubation on glass coverslips (top, glass) and LPS bilayers (bottom, LPS bilayer). Scale bar: 20 μm. (**b**) Plot showing the ratio of RelA fluorescence intensity in the cell nucleus to that in cytoplasm as a function of incubation time on various surfaces as indicated. Each data point represents mean□±□S.E. of results from a total of more than 500 cells in 10 areas from three independent experiments.

In resting cells, RelA was predominantly cytoplasmic, but upon stimulation with ligands, it translocated into the nucleus, which was stained with Hoechst dye (results after LPS stimulation shown in **Fig. 2a**). Different stimulation conditions resulted in varying percentages of cells displaying RelA translocation. In **Fig. 2b**, we present the nuclear-to-cytoplasmic fluorescence intensity ratio of RelA (Nuc/Cyt ratio) over time across three bilayer samples and two control samples, with the positive control being stimulation with 500 ng/mL soluble LPS and negative control being no stimulation on bare glass coverslips. In the positive control, cells exhibited rapid RelA translocation immediately after stimulation, confirming their inflammatory response. Stimulation with either the LPS+IgG bilayer or the LPS bilayer caused a gradual increase in RelA translocation, reaching a plateau approximately 60 min after initial stimulation. Notably, co-stimulation by the LPS+IgG bilayer resulted in faster and more pronounced RelA translocation compared to LPS alone. Intriguingly, cells stimulated on the IgG bilayer alone showed minimal RelA translocation, which we confirmed in numerous repeated experiments. Nevertheless, these results suggest that the observed increase in NF-κB activation was not solely due to the independent activation of two separate receptors, but rather a synergistic effect from their co-activation.

Because NF-κB activation drives cytokine production, we next quantified the effect of ligand stimulation on the secretion of the pro-inflammatory cytokine TNF-α. After stimulating cells on different surfaces for 6 hours, we measured TNF-α concentrations in the supernatants using enzyme-linked immunosorbent assay (ELISA) (**Fig. 3**). Cells co-stimulated on LPS+IgG bilayers secreted significantly higher levels of TNF-α compared to those stimulated on either the IgG bilayer or the LPS bilayer alone. These findings confirm that the co-activation of TLR4 and FcγR synergistically enhances inflammatory cytokine secretion.

**Fig. 3.**
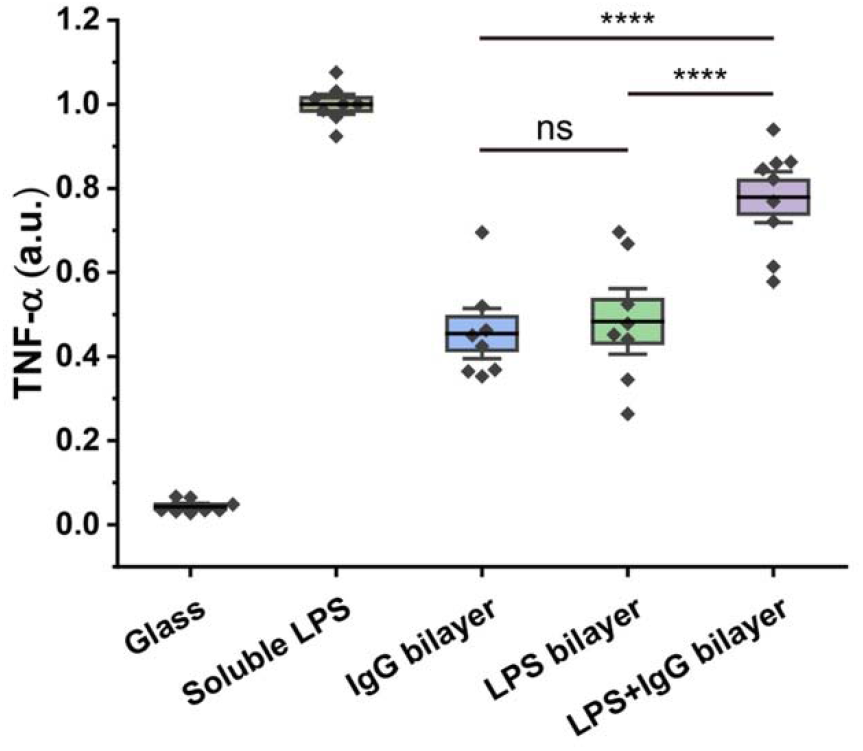
Measurements of cytokine secretion. ELISA analysis of TNF-α production in RAW264.7 macrophages incubation on various surfaces as indicated. Results were normalized against that from positive control (soluble LPS) and are presented as mean□±□SE. Data are representative of eight independent experiments each performed in four duplicates. Statistical significance is highlighted by p-values (Student’s t test) as follows: \*\*\*\**p*□≤ 0.0001; ns *p* > 0.05.

### Enhanced pY signaling from TLR4 and FcγR co-activation

Following the measurements of the inflammatory response, we examined how TLR4 and FcγR synergistic crosstalk affects early proximal signaling during receptor activation by quantifying phosphotyrosine (pY) levels, a key marker of receptor signaling for both FcγR and TLR4 ^15,22^. Macrophages were stimulated on various surfaces for 30 min, followed by pY immunostaining and imaging using TIRF microscopy (**Fig. 4a**).

**Fig. 4.**
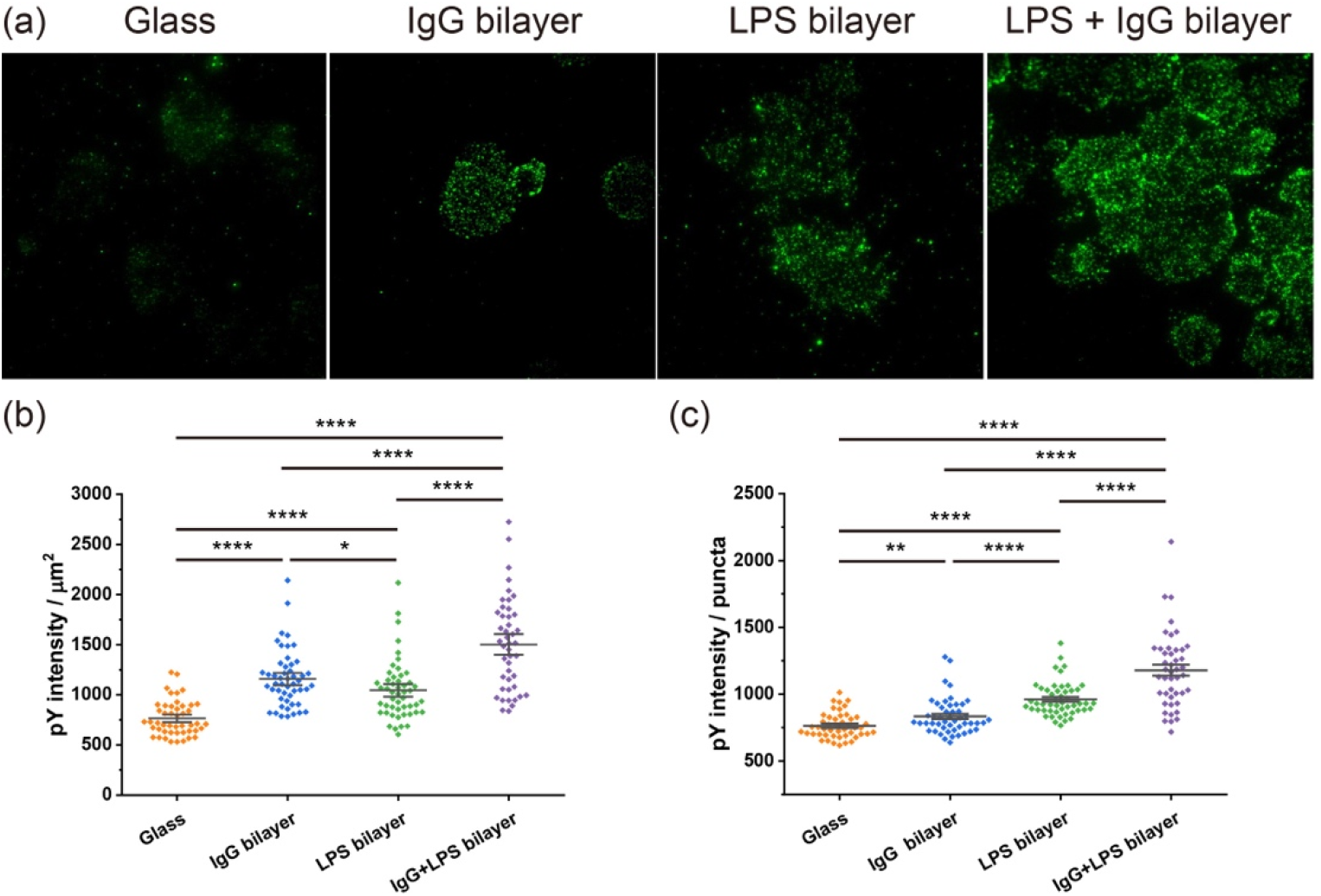
Measurements of phosphotyrosine (pY) levels. (**a**) Fluorescence images showing immunostained pY in RAW264.7 macrophage cells after 30 min of incubation on various substrates as indicated. Scale bar: 10 μm. (**b**) Quantification of pY fluorescence intensity of individual cells. For direct comparison between different cells, pY intensity of each cell was obtained as intensity per cell area. (**c**) Quantification of fluorescence intensity of individual pY puncta. Data are presented as mean ± SE obtained from 47 cells (control), 50 cells (LPS bilayer), 48 cells (IgG bilayer), and 45 cells (LPS+IgG bilayer) from three independent experiments. Statistical significance is highlighted by p values (Student’s t test) as follows: \*\*\*\**p* ≤ 0.0001; \*\**p* ≤ 0.01; \**p* ≤ 0.05.

Compared to cells on bare glass, those stimulated by ligands on bilayers exhibited significantly higher pY levels, confirming receptor activation. To quantitatively assess pY signaling at both the cellular and single receptor levels, we calculated pY intensity per μm^2^ and per puncta (**Fig. 4b, c**). In both metrics, cells stimulated on LPS+IgG bilayers displayed the highest pY intensities. These results are consistent with our findings on NF-κB activation and TNF-α secretion, indicating that the synergistic signaling initiated by the co-activation of TLR4 and FcγR extends from receptor engagement to downstream cytokine production.

### Synergistic effects of TLR4 and FcγR co-activation on receptor nanocluster

To further investigate how TLR4 and FcγR receptors interact at the cell membrane during synergistic signaling, we employed super-resolution direct stochastic optical reconstruction microscopy (dSTORM) to visualize the localization of individual TLR4 and FcγR receptors (**Fig. 5**). Cells were stimulated on various surfaces for 20 min before immunostaining. TLR4 was labeled with an anti-TLR4 primary antibody and a Cy3B-conjugated secondary antibody. Due to the challenge of specifically immunostaining FcγR, which binds to Fc regions of antibodies, we instead labeled phosphorylated spleen tyrosine kinase (pSyk) as a proxy for activated FcγRs. Syk is a cytoplasmic adaptor protein that associates with activated FcγRs, and its phosphorylation mediates downstream receptor signaling ^23^, so the localization of pSyk indicates activated FcγR. pSyk was labeled with an Alexa Fluor 647 (AF647)-conjugated anti-pSyk antibody. Reconstructed dSTORM images revealed that ligand-stimulated receptors formed nanoscale clusters.

**Fig. 5.**
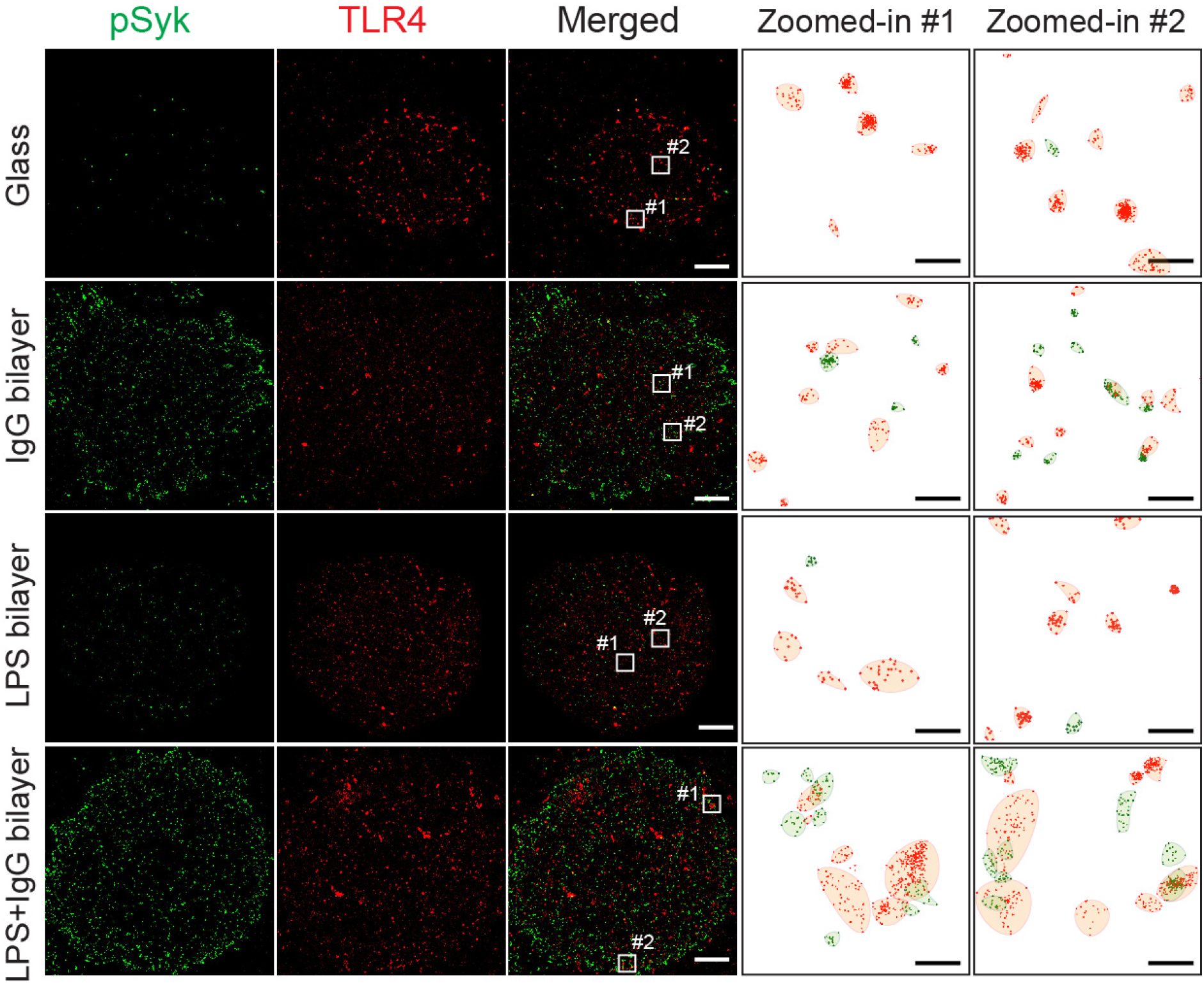
TLR4 and pSyk nanoclusters on various surfaces. Dual-color dSTORM images of TLR4 (red) and pSyk (green) in RAW264.7 macrophages on glass coverslips and ligand-conjugated lipid bilayers. Cluster maps of zoomed-in areas show the distribution of receptor nanoclusters identified using the topological mode analysis tool (ToMATo) method. Scale bars: 3 μm (dSTORM), 300 nm (zoomed-in).

To quantify receptor nanoclusters and their interactions, we used the topological mode analysis tool (ToMATo)^24^ after post-image processing that removed artifacts such as non-blinking dyes. The ToMATo method employs a persistence-based clustering algorithm to distinguish true clusters of single-molecule localizations from non-clustered events. Detailed parameter setup and validation for this analysis have been described in our previous work ^11^. Using this approach, we observed spatial overlap between TLR4 and FcγR nanoclusters upon co-activation (Fig. 5). Further analysis showed that co-activation of TLR4 and FcγRs significantly increased both the number and size of receptor clusters (**Fig. 6a, b**). When the receptors were activated individually, they each formed nanoclusters predominantly 50-70 nm in size, as determined by the peak of the size distribution. In contrast, co-activation resulted in larger TLR4 nanoclusters (∼120 nm), while pSyk nanoclusters, indicating activated FcγRs, also increased in size (∼76 nm).

**Fig. 6.**
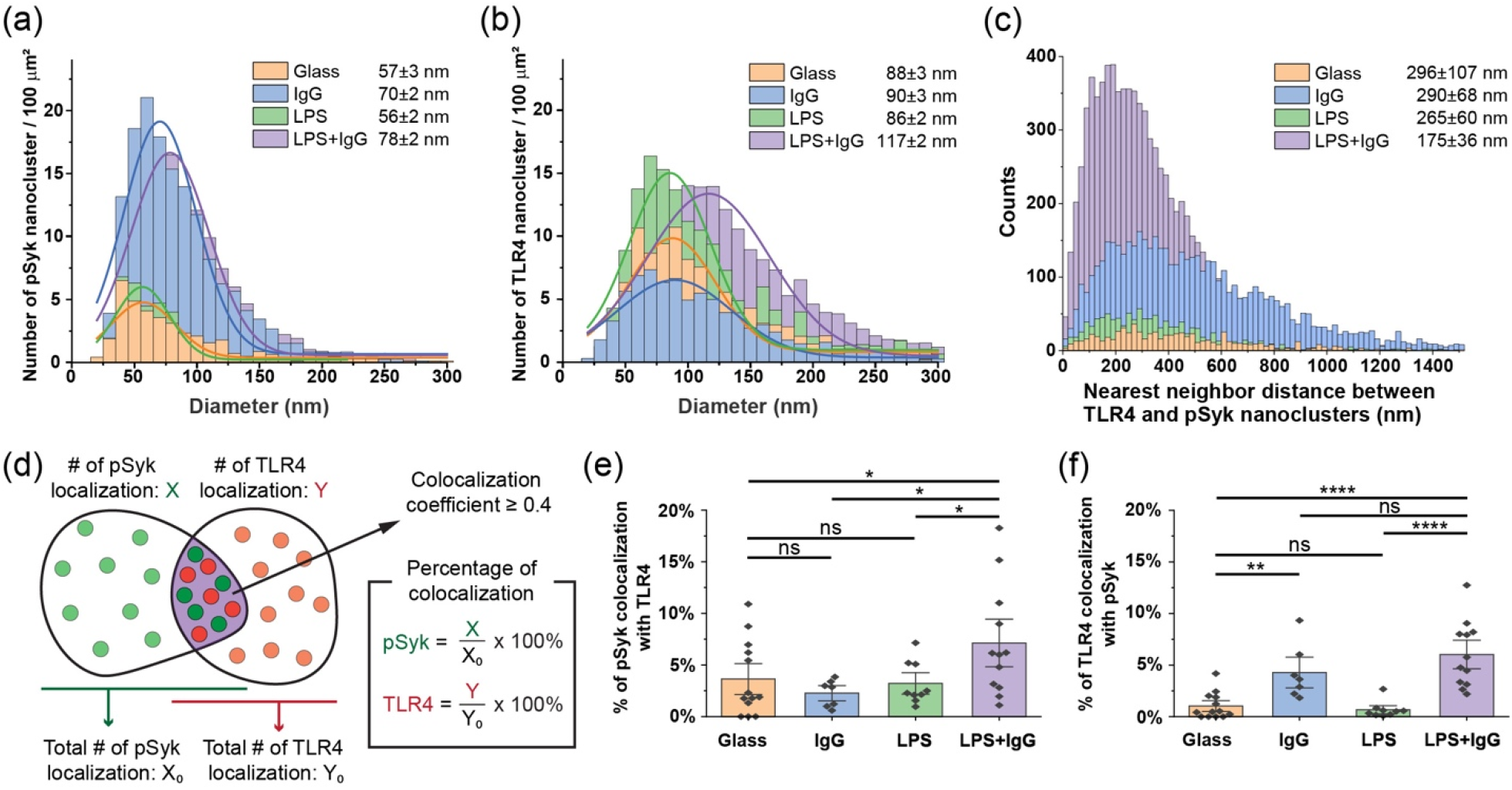
Quantification of TLR4 and pSyk nanoclusters. (**a, b**) Size distributions of pSyk and TLR4 nanoclusters in RAW264.7 cells on various surfaces. The legend in each graph indicates peak ± SE obtained from the histogram fitting. (**c**) Histograms showing the distribution of nearest neighbor distances measured from individual TLR4 nanoclusters to the closest pSyk nanoclusters. (**d**) Schematic illustration demonstrating the definition of the percentage of colocalization between nanoclusters. (**e, f**) The percentage of pSyk cluster colocalization with TLR4 and percentage of TLR4 cluster colocalization with pSyk. Data are presented as mean ± SE from 17 cells (control), 16 cells (IgG bilayer), 16 cells (LPS bilayer), and 16 cells (LPS+IgG bilayer) from four independent experiments. Statistical significance is highlighted by p values (Student’s t test) as follows: \*\*\*\**p* ≤ 0.0001; \*\**p* ≤ 0.01; \**p* ≤ 0.05; ns *p* > 0.05.

To quantify the partial overlap of nanoclusters, we first analyzed the centroid-to-centroid nearest neighbor distance (NND) between TLR4 and FcγR clusters upon co-activation (**Fig. 6c**). The NND was (290 ± 68) nm for FcγR activation on IgG bilayers and (265 ± 60) nm for TLR4 activation on LPS bilayers. However, co-activation on LPS+IgG bilayers reduced the NND to (175 ± 36) nm, which is smaller than the combined radii of TLR4 and pSyk clusters (195 ± 3 nm), suggesting that the clusters moved closer to each other, resulting in partial overlap during receptor crosstalk.

To directly quantify the spatial overlap between TLR4 and pSyk nanoclusters, we used a coordinate-based colocalization analysis ^25^ to calculate the percentage of TLR4 receptors colocalized with pSyk, and vice versa (**Fig. 6d-f**). Colocalization was determined by calculating the correlation coefficient between localizations, ranging from −1 (perfectly segregated) to +1 (perfectly colocalized), with values above 0.4 indicating colocalization, as previously validated^11^. The analysis revealed a significant increase in the percentage of colocalized TLR4 and pSyk upon co-activation, compared to the activation of either receptor alone (**Fig. 6e, f**). This supports the conclusion that receptor nanoclusters partially overlap during signaling crosstalk. The increased physical interaction likely results from the more and larger TLR4 and FcγR clusters formed during co-activation. These overlapping regions act as “signaling hotspots,” where TLR4 and FcγR, along with downstream signaling molecules, are brought into proximity to facilitate receptor crosstalk.

## Discussion

Previous studies have shown that TLR4 and FcγR function synergistically to regulate inflammatory responses in innate immune cells^7,9,10,13^, but the mechanism of their signaling crosstalk through receptor interactions remained unclear. In this study, we tested the hypothesis that this synergistic crosstalk in RAW264.7 macrophages is driven by the spatial organization of receptor nanoclusters. By using supported lipid bilayers functionalized with LPS and IgG, we achieved precise control over ligand presentation, including surface density, enabling us to draw quantitative conclusions about receptor interactions.

Our findings reveal that co-activation of TLR4 and FcγR significantly enhances both NF-κB activation and TNF-α secretion compared to activating either receptor alone. Importantly, super-resolution imaging showed that not only TLR4 and FcγR form discreet nanoclusters, but their co-activation leads to larger receptor nanoclusters and more interface overlapping between clusters, facilitating their physical interaction for synergistic signaling. This spatial proximity and increased cluster size promote enhanced tyrosine phosphorylation and amplify downstream inflammatory signaling.

An important finding is that this mechanism of receptor crosstalk through spatial reorganization is not unique to TLR4 and FcγR. We have previously shown that other receptor pairs, including Dectin-1 and TLR2, as well as FcγR and TLR2, also interact in similar ways during their signaling crosstalk^12,26^. This new evidence reinforces the general applicability of spatial clustering as a key mechanism for receptor signaling crosstalk. Previous studies have shown that FcγR and TLR2 co-precipitate in co-immunoprecipitation experiments, leading to the assumption that these receptors form heterodimers to mediate their synergistic effects^10^. Our study presents direct evidence challenging this assumption: although FcγR and TLR2 can form heterodimers within the overlapping regions of their nanoclusters, it is the nanoclusters themselves that function as the primary signaling units, facilitating their interactions and crosstalk. Altogether, our results highlight that spatial organization plays a crucial role in enhancing receptor crosstalk, leading to more effective immune responses through increased cytokine production and secretion. This underscores the importance of receptor nanoclusters as platforms for immune signaling amplification.

### Materials and methods Cells and reagents

RAW264.7 macrophage cells were purchased from ATCC (Manassas, VA) and cultured in Dulbecco’s modified Eagle medium (DMEM) supplemented with 10% fetal bovine serum, 100 U/mL penicillin, 100 mg/mL streptomycin, and 0.2 mM L-glutamine, all purchased from Thermo Fisher Scientific (Waltham, MA). RAW264.7 cells stably expressing EGFP-RelA were provided by Dr. Iain D. C. Fraser at National Institutes of Health. The cells were cultured in the DMEM without penicillin and streptomycin. The cells were maintained in an incubator at 37 °C with 5% CO_2_ and 95% relative humidity. Tumor necrosis factor (TNF)-α mouse uncoated ELISA Kits were purchased from Invitrogen (Waltham, MA).

Phospholipids, including 1,2-dioleoyl-sn-glycero-3-phosphocholine (DOPC), 1,2-dioleoyl-sn-glycero-3-phosphoethanolamine-N-(cap biotinyl) (sodium salt) (biotin-DOPE), and 1,2-dioleoyl-sn-glycero-3-phosphoethanolamine-N-(lissamine rhodamine B sulfonyl) (ammonium salt) (Rhodamine-DOPE), were purchased from Avanti Polar Lipids (Alabaster, AL). Reagents for ligand conjugation on lipid bilayers included streptavidin, NHS-Biotin, lipopolysaccharides from *Escherichia coli* (LPS), and Alexa Fluor™ 488 conjugated lipopolysaccharides from *Escherichia coli* Serotype 055:B5 (AF488-LPS), all purchased from ThermoFisher Scientific (Waltham, MA). NHS-CF640R was purchased from Biotium (Fremont, CA). Phosphate-buffered saline (PBS) (1×, pH 7.4), 2-amino-2-(hydroxymethyl)propane-1,3-diol (Tris), 4-(2-hydroxyethyl)-1-piperazineethanesulfonic acid (HEPES,), albumin from bovine serum (BSA), and immunoglobulin G (IgG) from rabbit serum, and Amicon Ultra filters (100K) were purchased from Sigma-Aldrich (St. Louis, MO). For biotinylation, 2 mg/mL IgG was conjugated with 0.3 μM NHS-biotin in 0.1 M sodium bicarbonate buffer (pH 8.25) at room temperature for 2 h on rotor. Excess of NHS-Biotin was removed by centrifugation using Amicon 100K filter 5 times with 14,000 rcf at 4°C for 10 min each. The same procedure was used for preparing fluorescent dye labeled CF640R-Biotin-IgG.

Reagents used for immunostaining included paraformaldehyde (PFA) and triton X-100, both from Sigma-Aldrich (St. Louis, MO), and Hoechst 33342 and anti-CD284 (TLR4) polyclonal antibody from ThermoFisher Scientific (Waltham, MA), and AF647-labeled anti-pSyk (Tyr525/526) rabbit monoclonal antibody from Cell Signaling Technology (Danvers, MA), and AF488-labeled anti-phospho-tyrosine (pY) monoclonal antibody (pY20) monoclonal mouse antibody from Santa Cruz Biotechnology (Dallas, TX). Secondary antibodies, sheep anti-goat antibody from Thermo Fisher Scientific (Waltham, MA), were labeled with NHS-Cy3B purchased from Cytiva (Marlborough, MA). The dye labeling efficiency was 2-3 dyes per antibody. Reagents used for dSTORM imaging were purchased from Sigma-Aldrich (St. Louis, MO) and included: Tris, Buffer A (10 mM Tris (pH 8.0) and 50 mM NaCl), Buffer B (50 mM Tris-HCI (pH 8.0), 10 mM NaCl, and 10% w/v glucose), 100× GLOX solution (3.4 mg/mL Catalase from bovine liver and 0.056 mg/mL glucose oxidase from Aspergillus niger in Buffer A), and 1 M mercaptoethylamine (MEA) in water (pH 8.0). TetraSpeck fluorescent microspheres (0.1 μm) were purchased from Thermo Fisher Scientific (Waltham, MA).

### Preparation of glass supported lipid bilayers

Lipid bilayers were composed of 2 mol% biotin-DOPE and varying mol% DOPC depending on the inclusion of LPS and fluorescent lipids. For LPS-containing lipid bilayers, an additional 1 wt% LPS was included. Fluorescent lipid bilayers were prepared by adding 0.05 mol% Rhodamine-DOPE. SLBs were prepared using the vesicle rupture method, involving several steps. First, lipids of desired compositions were mixed in chloroform to a total lipid mass of 1.0 mg and then dried in a round-bottom flask using a rotary evaporator to from a lipid film. It is important to note that LPS was added after the freeze-thaw step, rather than to the initial lipid mixture, as adding it earlier was found to prevent successful vesicle formation. Second, the dried lipid film was hydrated in 1 mL of 10 mM HEPES buffer (pH 7.4) containing 137 mM NaCl and then sonicated for 30 min at 40 ^°^C to produce small lipid vesicles. Third, the lipid suspension underwent three freeze-thaw cycles, alternating between -80 ^°^C freezer and 40 ^°^C warm water bath, to generate unilamellar vesicles. For making LPS-containing bilayers, LPS solution (1 mg/mL in Milli-Q water) was added to the lipid vesicle suspension at 1 wt% final concentration after this freeze-thaw step. Lastly, the lipid suspension was extruded eleven times through a polycarbonate filter of 100 nm pore size to create uniform vesicles, which were stored in -80 ^°^C freezer until use. The extrusion step was performed immediately before the experiment to ensure quality of SLBs. Prior to vesicle rupture, round glass coverslips were cleaned using a freshly prepared piranha solution (3:1 H_2_SO_4_:H_2_O_2_, v:v) for 20 min at room temperature, rinsed thoroughly using Milli-Q water, and then assembled into a custom-made imaging chamber. A final concentration of 0.1 mg/mL lipid vesicles in HEPES buffer was added to the chamber, and CaCl_2_ was included at a final concentration of 2 mM to enhance SLBs formation on the glass surface. The SLBs were allowed to form for 1 h at 37 ^°^C before being rinsed with 1×PBS and then blocked with BSA solution (0.01 w/v% in PBS) for 45 min.

### Functionalization of ligands on lipid bilayers

To prepare IgG-functionalized lipid bilayers, bilayers were first incubated with 20 μg/mL streptavidin in 1×PBS containing 1% (w/v) BSA for 1 h at room temperature. After washing away excess streptavidin, bilayers were incubated with 20 μg/mL IgG-biotin in 1×PBS containing 1% (w/v) BSA for 1 h at room temperature. The functionalized SLBs were extensively rinsed with PBS and used immediately for cell experiments. The same procedure was used to prepare bilayers containing only IgG or both IgG and LPS.

To quantify the surface density of IgG on SLBs, non-fluorescent IgG-biotin was mixed with CF640R dye labeled IgG-biotin at a molar ratio of 10,000:1 in 1× PBS. The mixed ligands were conjugated to the bilayers using the same procedure described above. Similarly, to quantify the surface density of LPS on bilayers, a mixture of non-fluorescent LPS and AF488-LPS at a molar ratio of 10,000:1 was added during lipid bilayer preparation. The bilayers were imaged using total internal reflection fluorescence (TIRF) microscopy. The total number of fluorescent IgG or LPS in a 40×40 μm region of each given image was counted using the TrackMate plugin of ImageJ software. Surface density of IgG or LPS was calculated by multiplying the surface densities of fluorescent IgG or LPS with the mixing molar ratio (10,000 for IgG and 10,000 for LPS).

### Immunofluorescence staining

RAW264.7 macrophage cells were seeded on lipid bilayers at a density of approximately 0.5×10^6^ cells/mL. After a 20 min incubation at 37 °C with 5% COL, the cells were washed with 1× PBS, fixed with 4% (w/v) PFA in 1× PBS for 30 min at room temperature, and then rinsed three times with 1× PBS for 5 min each. The cells were permeabilized with Triton X-100 for 30 min at room temperature, followed by three more rinses with 1× PBS for 5 min each. The cells were then passivated in a blocking buffer containing 0.1% (v/v) Tween 20, 5% (w/v) BSA, and 22.52 mg/mL glycine in 1× PBS for 1 h at room temperature. For immunostaining of pSyk and TLR4, the cells were first incubated with AF647-labeled anti-pSyk primary antibodies (5 μg/mL) in the presence of 5% (w/v) BSA for 1 h at room temperature. After this, the cells were incubated with anti-TLR4 goat IgG (10 μg/mL) for 1 h, followed by incubation with Cy3B-labeled sheep anti-goat secondary antibody (2 μg/mL) in the presence of 5% (w/v) BSA for an additional 1 h.

For immunostaining of pY, cells were incubated on lipid bilayers or bare glass surfaces for 30 min. After following the same fixation and permeabilization steps described above, cells were incubated with AF488-labeled anti-pY (5 μg/mL) in the presence of 5% (w/v) BSA for 1 h at room temperature. Each antibody incubation was followed by three washes with washing buffer (0.1% Tween 20 and 1% BSA in 1× PBS) for 15 min.

### Epifluorescence and total internal reflection fluorescence (TIRF) Microscopy

Epifluorescence and TIRF images were acquired using a Nikon Eclipse-Ti inverted microscope, equipped with a Nikon 100×/1.49 N.A. oil-immersion TIRF objective or a Nikon 40×/0.95 N.A. air-immersion objective and a Hamamatsu ORCA-Fusion digital CMOS camera.

### Analysis of fluorescence recovery after photobleaching (FRAP)

FRAP measurements were performed at 37 ^°^C using epifluorescence microscopy. A circular region of 39.5 μm in diameter was photobleached for 30 s, and post-bleach fluorescence images were acquired at a rate of 2 s/frame. To identify the area for FRAP analysis, the fluorescence intensity was first normalized to that of an unbleached area to correct for additional photobleaching. The intensity versus time data during recovery was plotted and fitted with a two-phase exponential association curve using OriginPro software (OriginLab). Diffusion coefficient (*D*) was calculated using the following equation^27^:

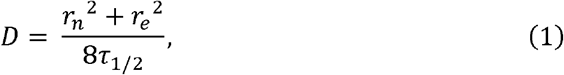

Where τ_1/2_ is the half recovery time of fluorescence intensity, *r*_*n*_= 19.75 μm is the radius of photobleached area, and *r*_*e*_ is the effective radius obtained by fitting the post-bleach profile using the equation:

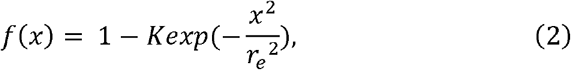

In Eq. 2, *f* (*x*) is the fluorescence intensity at distance *x* along the diagonal of a post-bleached image, and *K* is a fitting parameter that represents the bleaching depth. The initial value of *K* was set to 1. The same procedure was used to measure mobility of lipid bilayers containing AF488-LPS.

### Quantification of NF-κB RelA nuclear translocation

RAW264.7 macrophages stably expressing EGFP-RelA were plated on glass coverslips or lipid bilayers at a density of approximately 0.5×10^6^ cells/mL and incubated for various durations as specified. As positive control, 500 ng/mL LPS solution was added to the cell culture medium during incubation. Following incubation, cells were fixed for 10 min with 4% (w/v) PFA in 1× PBS and stained with Hoechst 33342 (0.1 μg/mL) in 1× PBS for 10 min at room temperature. To assess the nuclear translocation of NF-κB RelA, the fluorescence intensities of EGFP-RelA within the cell nucleus and cytoplasm were measured (**Fig. S1**). The nuclear-to-cytoplasmic intensity ratio of EGFP-RelA was quantified using the following method^18^. Briefly, original images in each channel, Nucleus and EGFP-RelA, (**Fig. S1b**) were converted into binary images using automatic local thresholding function in ImageJ (**Fig. S1c**). Here, the nucleus mask (**Fig. S1d, upper raw**) and cytoplasmic mask (**Fig. S1d, lower raw**) defined the region of interest (ROI) for the cell nucleus and cytoplasm, respectively. The cytoplasmic mask was created by subtracting the nucleus mask from the cell mask (**Fig. S1c, lower raw**). The resulting ROI masks (**Fig. S1d**) were then applied to the original EGFP-RelA image (**Fig. S1b, lower raw**) to generate separated EGFP-RelA images on nuclear and cytoplasmic regions (**Fig. S1e**). Fluorescence intensities per pixel for both the nuclear and cytoplasmic regions were measured using ImageJ software. The nuclear-to-cytoplasmic fluorescence intensity ratio of EGFP-RelA was calculated to quantify the extent of NF-κB RelA nuclear translocation.

### Cytokine assays

RAW 264.7 macrophages cells were incubated on glass coverslips or lipid bilayers at a density of approximately 0.5×10^6^ cells/mL for 6 h at 37 ^°^C with 5% CO_2_ and 95% relative humidity. Concentration of TNF-α secreted into the supernatants was quantified using ELISA kits according to the manufacturer’s protocol. For the positive control, 500 ng/mL LPS was added to the cell culture medium during incubation. Absorbance was recorded at 450 nm using a BioTek Synergy H1 microplate reader. In each sample, four duplicates were measured, and three independent experiments were performed for each condition.

### Analysis of pY activation and signaling

To measure the fluorescence intensity of individual pY puncta, a single-particle localization algorithm was used to determine the centroid and the full width at half-maximum of the point-spread function for each pY punctum’s fluorescence intensity profile^28^. The analysis provided the number of pixels (pY signals) and the integrated intensity of each punctum. To measure the pY intensity in each cell, a mask was created based on pY intensity threshold to distinguish individual cells (**Fig. S2**). The pY intensity per unit area within the mask was calculated by dividing the integrated pY intensity divided by the total number of pixels in the mask. The fluorescence intensity of a single punctum was calculated as the intensity per pixel, calculated by dividing the integrated intensity by the number of pixels of the punctum.

### Image acquisition of Direct Stochastic Optical Reconstruction Microscopy (dSTORM)

All dSTORM imaging was performed in TIRF mode using a Nikon Ti2 Eclipse microscope equipped with a perfect focus system, a TIRF ×100 oil-immersion objective (NA 1.49), and an Andor iXon3 EMCCD camera. Dual-color dSTORM images were acquired sequentially with the excitation of 637 and 561 nm diode lasers at power densities of 10 kW/cm^2^ and 8 kW/cm^2^, respectively. Each fluorescence channel was imaged with an exposure time of 10 ms for a total of 10,000 consecutive frames. A 405 nm diode laser at a power density of 0.5 kW/cm^2^ was used to activate fluorophores stochastically. Fixed and immunostained cell samples were maintained in freshly prepared dSTORM imaging buffer, consisting of 100 mM MEA and 1× GLOX solution in Buffer B, and imaged at room temperature. For each sample, the dSTORM imaging buffer was freshly prepared and replaced immediately prior to the image acquisition to ensure optimal dye blinking.

### Post-processing of dSTORM images

#### 1. Image reconstruction

dSTORM images were reconstructed using ThunderSTORM plugin in ImageJ software^29^. Briefly, the centroids of activated molecules were localized in each frame by fitting their point-spread functions (PSF) using least-square Gaussian fitting method. To resolve individual molecules in area of high spatial densities, a multiple emitter fitting analysis was employed^30^. Sequential filters were applied to eliminate noise from the images.

(1) An intensity filter with a threshold of *N* > α*b* was applied to eliminate molecules with low photon counts. Here, *N* represents the number of photons detected from a given molecule, *b* is the standard deviation of the signal within the fitting region, and α, a sensitivity factor typically ranging from 20 to 40, was set as 30 in this study^31^.
(2) To eliminate noise from nonclustered molecules, a local density filter was applied. For each localization, the number of neighboring molecules within a specific radius was counted, and if this number fell below a set threshold, that localization was discarded. In this study, a threshold of at least 10 neighbors within a 100 nm radius was used to effectively filter out the nonclustered molecules.
(3) To eliminate multiple blinking from a single molecule across sequential frames, localizations detected within a single pixel of the CCD camera (105 nm in this study) over five consecutive frames were treated as same localization. They were merged into a single localization by averaging the coordinates from all involved localizations. The photon count and background signal level for the new, merged localization were calculated by summing these parameters from the pre-merged localizations. The inaccuracy of the newly merged localization was calculated using following as:

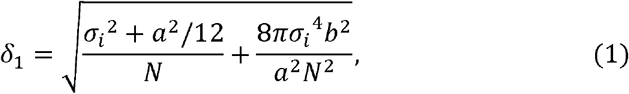

where *N* is the number of photons collected from a given molecule, σ_*i*_ is the fitted standard deviation of PSF of the detected molecules in either the *x* or *y* direction, *a* is the pixel size of the CCD camera, and *b* is the standard deviation of background.
(4) To eliminate repeated localizations of a single molecule within the same frame, such as when multiple dyes on a single protein blink simultaneously and are detected, localizations within the scale of the localization inaccuracy were grouped together. Within each group, only the molecule with the smallest localization inaccuracy was retained.

#### 2. Registration and correction of multicolor fluorescence dSTORM images

For dual-color dSTORM images, the images were processed separately for each channel, and the lateral drift between two fluorescence channels was corrected using TetraSpeck fluorescent particles (100 nm in diameter) as fiducial markers. These markers were added to the cell samples right before dSTORM imaging. Since dual-color dSTORM imaging was performed sequentially–first with the 637 nm excitation channel, followed by the 561 nm channel–the first frame of the 637 nm channel was used as reference to correct the lateral drift of both channels. Briefly, the *x*–*y* coordinates of each fiducial marker (*x*_*m*, *N*_ and *y*_*m*, *N*_) in each frame of the 637 nm channel were averaged to obtain the following:

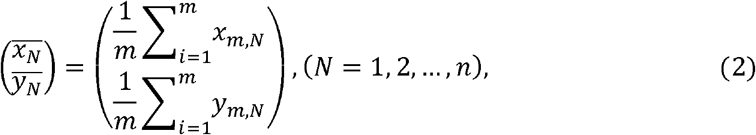

where *m* is the number of fiducial markers in each frame and *N* is the total frame number. The averaged coordinates of the fiducial markers 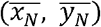 in each frame *n* were then compared to those in frame 1 to obtain the average displacement of each frame relative to frame 1:

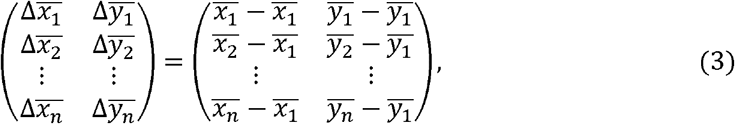

The relative averaged displacement 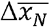 and 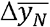 in each frame *n* were then subtracted from every single-molecule localization *X*_*n*_ and *Y*_*n*_ in the corresponding dSTORM image of the cell sample, generating the corrected *x*–*y* localization 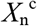 and 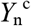 in each frame:

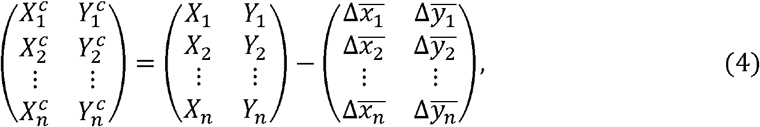

where *X*_*n*_ and *Y*_*n*_ represent all the original *x* and *y* coordinates of the cell sample in each frame *n*. 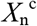 and 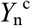 represent all the corrected *x*–*y* coordinates from each frame *n*. The same procedure was applied to correct lateral drift in images from the 561 nm excitation channel using frame 1 in 637 nm channel as a reference. The final dSTORM images were visualized using either averaged shifted histogram^32^ or single-molecule localization scatter plot. Averaged shifted histogram was achieved using ThunderSTORM plugin in ImageJ with a standard deviation equal to the localization inaccuracy. Single-molecule localization scatter plot was generated using the R package ‘ggplot2’^33^.

#### 3. Identification of receptor nanocluster

After post-processing of dSTORM images, single-molecule localizations of TLR4 and pSyk were grouped into nanoclusters using the Topological Mode Analysis Tool (ToMATo)^24^, a persistence-based clustering segmentation algorithm^34^. In essence, this algorithm first calculates the detection density at each single-molecule localization by counting the number of neighboring detections within a circle of fixed search radius *r*. Candidate clusters are then formed using a mode-seeking approach, where each single-molecule localization is linked to neighboring localizations with the highest detection density within the search radius, *r*. The linked localizations collectively form a candidate cluster. Each candidate cluster is characterized by a point of maximum density (birth density) and a saddle point (death density). The difference between the birth and death densities is defined as the persistence of the candidate cluster. To refine the clustering, a persistence threshold is determined using a persistence diagram, which plots the birth and death densities of each candidate cluster on a Cartesian coordinate system. The persistence threshold, τ, is defined as the x-axis value at the intersection of the line *y* = *x* − τ with the birth density. Candidate clusters with persistence below this threshold are merged with neighboring clusters that meet or exceed the threshold.

In this study, we set a search radius *r* = 50 and a persistence threshold τ = 5, following a previously reported simulation that evaluated ToMATo’s performance across various parameters^24^. After identifying the clusters, the ToMATo clustering analysis was executed using the R package “RSMLM”^24^. The cluster area was defined as the convex hull encompassing all single-molecule localizations within a cluster. In our study, we identified a mode value of 10 localizations per receptor nanocluster (both TLR4 and pSyk). Consequently, we used a threshold of “localizations ≥ 10” as the minimum number of localizations per receptor nanocluster, aiming to minimize the likelihood of recognizing non-specifically adsorbed antibodies for genuine receptor nanoclusters.

#### 4. Analysis of receptor nanocluster overlap

The degree of overlap between two types of receptor nanoclusters was determined using a coordinate-based colocalization (CBC) analysis^25^, which is available through ThunderSTORM plugin in ImageJ. In brief, for each single-molecule localization (*A*_*i*_) of protein A, the density of nearby single-molecule localizations of both protein A and protein B within a circle of radius *r* around *A*_*i*_ was calculated using

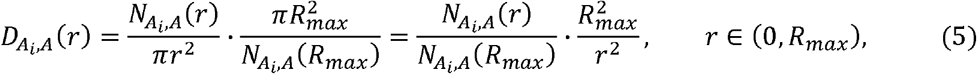

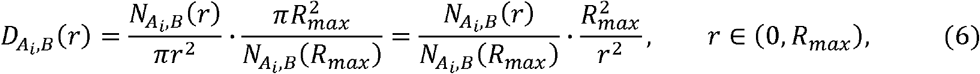

where 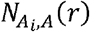 and 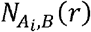 represent the number of localizations of protein A and B, respectively, within the the radius *r* surrounding *A*_*i*_. The maximum radius, *R*_*max*_, is set by the user. The density gradients 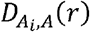 and 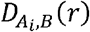 were then compared by calculating their Spearman correlation coefficient 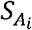 The colocalization coefficient (DoC) for each localization, 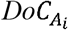 was computed as follows:

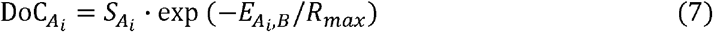

Where 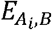 is the distance from *A*_*i*_ to its nearest neighbor protein *B* localization. For each single-molecule localization in the dual-color dSTORM images, a DoC score was obtained, ranging from −1 (anti-colocalized) to +1 (perfectly colocalized), with 0 indicating no colocalization. In this study, the analysis was performed on the single-molecule localizations of TLR4 and pSyk across increasing radii, using the following parameters: a radius step of 10 nm and a step count of 25. Single-molecule localizations with DoC scores of 0.4 or higher were considered colocalized. The overall degree of overlap was calculated by dividing the total number of colocalized localizations by the total number of localizations for each protein’s species.

## Data availability

The authors declare that all data supporting the findings of this study are available within the article and its Supplementary Information files.

## Acknowledgements

We thank Dr. Miao Li for providing R script for dSTORM post-processing and Dr. Iain D. C. Fraser at National Institutes of Health for providing the RAW264.7 cells expressing GFP-RelA. This work was supported by the National Institutes of Health (NIH) under award R35GM124918. The content is solely the responsibility of the authors and does not necessarily represent the official views of the National Institutes of Health.

## Author contributions

S.L. performed experiments and analyzed data. Y.Y. designed research. S.L. and Y.Y. wrote the paper.

## Competing interests

The author(s) declare no competing interests.

## Additional Information

Supplementary information.

## Notes

### Competing Interest Statement

The authors have declared no competing interest.

## References

1 Lee, W. L., Harrison, R. E. & Grinstein, S. Phagocytosis by neutrophils. Microbes and Infection 5, 1299–1306 (2003). 10.1016/j.micinf.2003.09.014

2 Aderem, A. & Underhill, D. M. Mechanisms of phagocytosis in macrophages. Annual Review of Immunology 17, 593–623 (1999). 10.1146/annurev.immunol.17.1.593

3 Iwasaki, A. & Medzhitov, R. Control of adaptive immunity by the innate immune system. Nature Immunology 16, 343–353 (2015). 10.1038/ni.3123

4 Uribe-Querol, E. & Rosales, C. Phagocytosis: Our Current Understanding of a Universal Biological Process. Frontiers in Immunology 11 (2020). 10.3389/fimmu.2020.01066

5 Vogelpoel, L. T. et al. Fc gamma receptor-TLR cross-talk elicits pro-inflammatory cytokine production by human M2 macrophages. Nat Commun 5, 5444 (2014). 10.1038/ncomms6444

6 Hunt, D., Drake, L. A. & Drake, J. R. Murine macrophage TLR2-FcgammaR synergy via FcgammaR licensing of IL-6 cytokine mRNA ribosome binding and translation. PLoS One 13, e0200764 (2018). 10.1371/journal.pone.0200764

7 Wenink, M. H. et al. The inhibitory Fc gamma IIb receptor dampens TLR4-mediated immune responses and is selectively up-regulated on dendritic cells from rheumatoid arthritis patients with quiescent disease. J Immunol 183, 4509–4520 (2009). 10.4049/jimmunol.0900153

8 Negishi, H. et al. Cross-interference of RLR and TLR signaling pathways modulates antibacterial T cell responses. Nature Immunology 13, 659-+ (2012). 10.1038/ni.2307

9 Shang, L. M. et al. Selective Antibody Intervention of Toll-like Receptor 4 Activation through Fc γ Receptor Tethering. Journal of Biological Chemistry 289, 15309–15318 (2014). 10.1074/jbc.M113.537936

10 Rittirsch, D. et al. Cross-talk between TLR4 and FcgammaReceptorIII (CD16) pathways. PLoS Pathog 5, e1000464 (2009). 10.1371/journal.ppat.1000464

11 Li, M., Vultorius, C., Bethi, M. & Yu, Y. Spatial Organization of Dectin-1 and TLR2 during Synergistic Crosstalk Revealed by Super-resolution Imaging. J Phys Chem B 126, 5781–5792 (2022). 10.1021/acs.jpcb.2c03557

12 Li, M. et al. Immobile ligands enhance FcgammaR-TLR2/1 crosstalk by promoting interface overlap of receptor clusters. Biophys J 121, 966–976 (2022). 10.1016/j.bpj.2022.02.010

13 Vargas-Hernandez, O. et al. THP-1 cells increase TNF-alpha production upon LPSL+Lsoluble human IgG co-stimulation supporting evidence for TLR4 and Fcgamma receptors crosstalk. Cell Immunol 355, 104146 (2020). 10.1016/j.cellimm.2020.104146

14 Lu, Y. C., Yeh, W. C. & Ohashi, P. S. LPS/TLR4 signal transduction pathway. Cytokine 42, 145–151 (2008). 10.1016/j.cyto.2008.01.006

15 Bournazos, S., Gupta, A. & Ravetch, J. V. The role of IgG Fc receptors in antibody-dependent enhancement. Nat Rev Immunol 20, 633–643 (2020). 10.1038/s41577-020-00410-0

16 Verma, A. et al. Analysis of the Fc gamma receptor-dependent component of neutralization measured by anthrax toxin neutralization assays. Clin Vaccine Immunol 16, 1405–1412 (2009). 10.1128/CVI.00194-09

17 Park, M. H. & Hong, J. T. Roles of NF-kappaB in Cancer and Inflammatory Diseases and Their Therapeutic Approaches. Cells 5 (2016). 10.3390/cells5020015

18 Noursadeghi, M. et al. Quantitative imaging assay for NF-kappaB nuclear translocation in primary human macrophages. J Immunol Methods 329, 194–200 (2008). 10.1016/j.jim.2007.10.015

19 Christian, F., Smith, E. L. & Carmody, R. J. The Regulation of NF-kappaB Subunits by Phosphorylation. Cells 5 (2016). 10.3390/cells5010012

20 Liu, T., Zhang, L., Joo, D. & Sun, S. C. NF-kappaB signaling in inflammation. Signal Transduct Target Ther 2, 17023-(2017). 10.1038/sigtrans.2017.23

21 Sun, S. C. Non-canonical NF-kappaB signaling pathway. Cell Res 21, 71–85 (2011). 10.1038/cr.2010.177

22 Chattopadhyay, S. & Sen, G. C. Tyrosine phosphorylation in Toll-like receptor signaling. Cytokine Growth Factor Rev 25, 533–541 (2014). 10.1016/j.cytogfr.2014.06.002

23 Kiefer, F. et al. The Syk protein tyrosine kinase is essential for Fcgamma receptor signaling in macrophages and neutrophils. Mol Cell Biol 18, 4209–4220 (1998). 10.1128/MCB.18.7.4209

24 Pike, J. A. et al. Topological data analysis quantifies biological nano-structure from single molecule localization microscopy. Bioinformatics 36, 1614–1621 (2020). 10.1093/bioinformatics/btz788

25 Malkusch, S. et al. Coordinate-based colocalization analysis of single-molecule localization microscopy data. Histochem Cell Biol 137, 1–10 (2012). 10.1007/s00418-011-0880-5

26 Li, W. Q., Yan, J. & Yu, Y. Geometrical reorganization of Dectin-1 and TLR2 on single phagosomes alters their synergistic immune signaling. P Natl Acad Sci USA 116, 25106–25114 (2019). 10.1073/pnas.1909870116

27 Kang, M., Day, C. A., Kenworthy, A. K. & DiBenedetto, E. Simplified equation to extract diffusion coefficients from confocal FRAP data. Traffic 13, 1589–1600 (2012). 10.1111/tra.12008

28 Parthasarathy, R. Rapid, accurate particle tracking by calculation of radial symmetry centers. Nat Methods 9, 724–726 (2012). 10.1038/nmeth.2071

29 Ovesny, M., Krizek, P., Borkovec, J., Svindrych, Z. & Hagen, G. M. ThunderSTORM: a comprehensive ImageJ plug-in for PALM and STORM data analysis and super-resolution imaging. Bioinformatics 30, 2389–2390 (2014). 10.1093/bioinformatics/btu202

30 Huang, F., Schwartz, S. L., Byars, J. M. & Lidke, K. A. Simultaneous multiple-emitter fitting for single molecule super-resolution imaging. Biomed Opt Express 2, 1377–1393 (2011). 10.1364/BOE.2.001377

31 Krizek, P., Raska, I. & Hagen, G. M. Minimizing detection errors in single molecule localization microscopy. Opt Express 19, 3226–3235 (2011). 10.1364/OE.19.003226

32 Scott, D. W. Averaged Shifted Histograms - Effective Nonparametric Density Estimators in Several Dimensions. Ann Stat 13, 1024–1040 (1985). 10.1214/aos/1176349654

33 Villanueva, R. A. M. & Chen, Z. J. ggplot2: Elegant Graphics for Data Analysis, 2nd edition. Meas-Interdiscip Res 17, 160–167 (2019). 10.1080/15366367.2019.1565254

34 Chazal, F., Guibas, L. J., Oudot, S. Y. & Skraba, P. Persistence-Based Clustering in Riemannian Manifolds. J Acm 60 (2013). 10.1145/2535927

