## Supplementary Figures 1 and 2 for "Nanocluster-Mediated Signaling Crosstalk between FcγR and TLR4 in Macrophage Inflammatory Responses"

**This PDF file includes:**

SI Figures S1 and S2

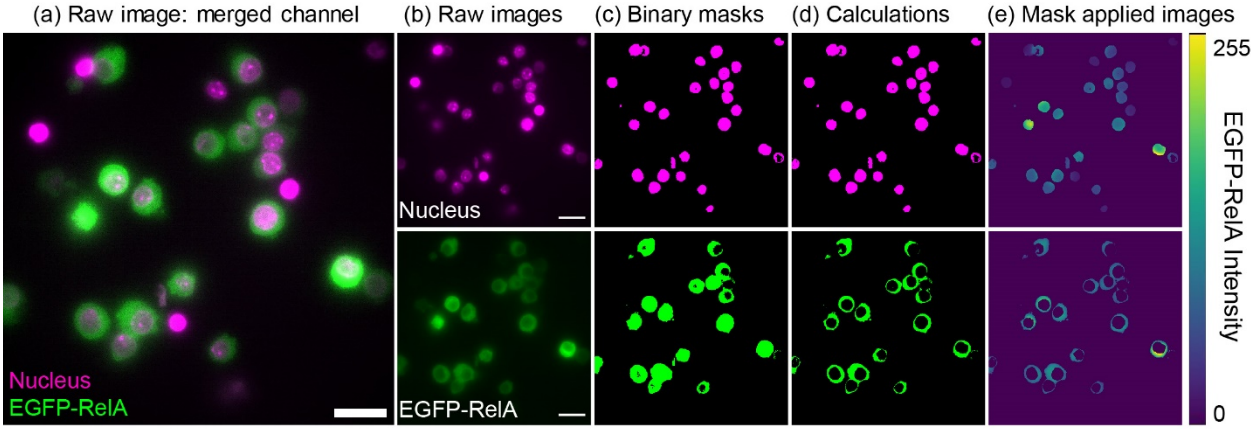


**Fig. S1. Quantification of NF-κB RelA nuclear translocation.** (a) Fluorescence image of EGFP-RelA expressing RAW264.7 (green) cells stained with Hoechst 33342 (magenta). Scale bar: 20 µm. (b) The original fluorescent image of nucleus (upper, magenta) and cytoplasm (lower, green) of EGFP-RelA expressing RAW264.7 cells. (c) Fluorescence images were converted to binary images through automatic local thresholding using ImageJ software. (d) Images of Hoechst 33342 were used as the nuclear mask; the cytoplasmic mask was then obtained by subtracting the nuclear mask from the cell mask. (e) The nuclear and cytoplasmic masks, respectively, were applied to the original EGFP-RelA images, to separately obtain the fluorescence intensity of RelA in cell nucleus and cytosol.

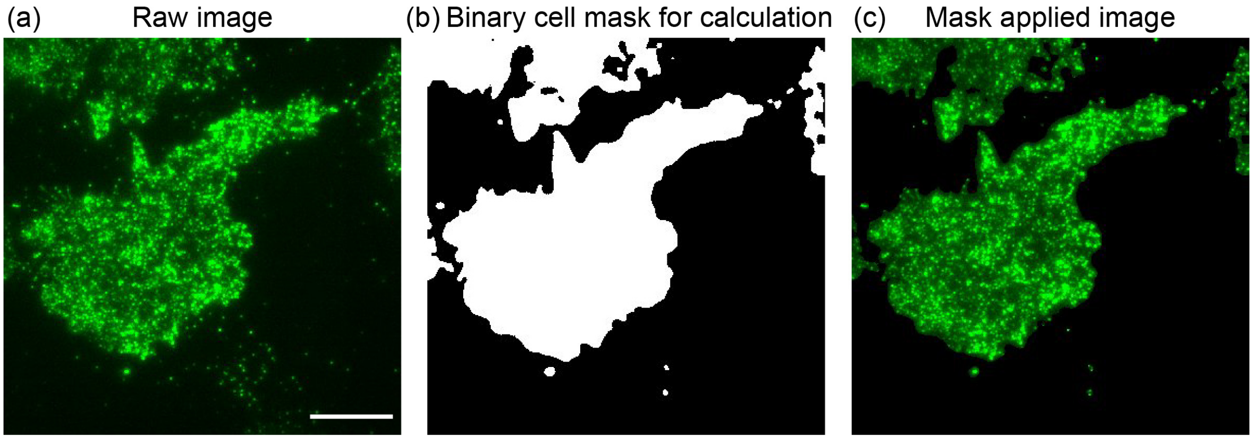


**Fig. S2. Quantification of pY fluorescence intensity.** (a) Original fluorescence image of RAW264.7 cells immunostained with Alexa488 labeled pY antibody. Scale bar: 10 μm. (b) The original fluorescence image (a) was converted to a binary image through automatic local thresholding using ImageJ software. (c) The binary mask was then applied to the original image to quantify the pY fluorescence signal per area.
